# High-fidelity *in vitro* packaging of diverse synthetic cargo into encapsulin protein cages

**DOI:** 10.1101/2024.11.24.625105

**Authors:** Taylor N. Szyszka, Rezwan Siddiquee, Alex Loustau, Lachlan S. R. Adamson, Claire Rennie, Tiancheng Huang, Reginald Young, Andrew Care, Yu Heng Lau

**Affiliations:** School of Chemistry, The University of Sydney, Camperdown, NSW 2006, Australia; The University of Sydney Nano Institute, The University of Sydney, Camperdown, NSW 2006, Australia; ARC Centre of Excellence in Synthetic Biology, The University of Sydney, Camperdown, NSW 2006, Australia; School of Life Sciences, University of Technology Sydney, Sydney, NSW 2007, Australia; Australian Institute for Microbiology and Infection, Sydney, NSW 2007, Australia; ARC Centre of Excellence for Innovations in Peptide and Protein Science, The University of Sydney, Camperdown, NSW 2006, Australia

## Abstract

Cargo-filled protein cages are powerful tools in biotechnology with demonstrated potential as catalytic nanoreactors and vehicles for targeted drug delivery. While endogenous biomolecules can be packaged into protein cages during their expression and self-assembly inside cells, synthetic cargo molecules are typically incompatible with live cells and must be packaged *in vitro*. Here we report a fusion-based *in vitro* assembly method for packaging diverse synthetic cargo into encapsulin protein cages that outperforms standard *in cellulo* assembly, producing cages with superior uniformity and thermal stability. Fluorescent dyes, proteins and cytotoxic drug molecules can all be selectively packaged with high efficiency *via* a peptide-mediated targeting process. The exceptional fidelity and broad compatibility of our *in vitro* assembly platform enables generalisable access to cargo-filled protein cages that host novel synthetic functionality for diverse biotechnological applications.

## Introduction

Cages constructed from self-assembling proteins are ubiquitous in nature, providing a rich source of engineerable molecular scaffolds for biotechnological applications.^1,2^ The most common class of engineered cages are virus-like particles (VLPs), formed when viral capsid proteins are assembled into hollow compartments in the absence of the viral genome.^3–5^ Other commonly engineered cages include iron-storing ferritins,^6,7^ *de novo* designed cages,^8^ cage-forming enzymes such as lumazine synthase,^9^ and prokaryotic proteinaceous organelles that include bacterial microcompartments^10,11^ and encapsulin nanocompartments.^12,13^ Each class of cage-forming proteins has distinct properties – e.g. size, thermal and chemical stability, tolerance to sequence and cargo engineering, immunogenicity – determining their suitability across diverse applications that include targeted delivery vehicles for synthetic drug molecules, vaccine scaffolds for antigen display, and catalytic nanoreactors for biomanufacturing.

The ability to package synthetic cargo into protein cages is a crucial limitation for many biotechnological applications. Protein cages typically assemble during their expression inside a cellular host, thus limiting cargo packaging to biomolecules that can be produced inside cells (Figure 1a). Indeed, we and many others have shown that protein cages can be readily reprogrammed to package cargo proteins or nucleic acids during assembly inside cells.^14–21^ To load synthetic non-biological cargo however, protein cages are often purified from their cellular hosts, disassembled under buffer conditions that disfavour assembly (e.g. changing pH, ionic strength, temperature, or adding chemical denaturants), then reassembled in the presence of the cargo.^22–31^ Drawbacks of this approach include the risk of damage to cage proteins or cargo during exposure to potentially harsh disassembly conditions, as well as suboptimal cargo loading efficiency when a passive statistical encapsulation mechanism is used.^32–38^ Furthermore, the subsequent *in vitro* reassembly process can have poor fidelity, generating a significant proportion of defective or aggregation-prone cages with inferior structural integrity, uniformity and thermal stability relative to their original correctly-assembled native state.^22,23,39,40^

**Figure 1.**
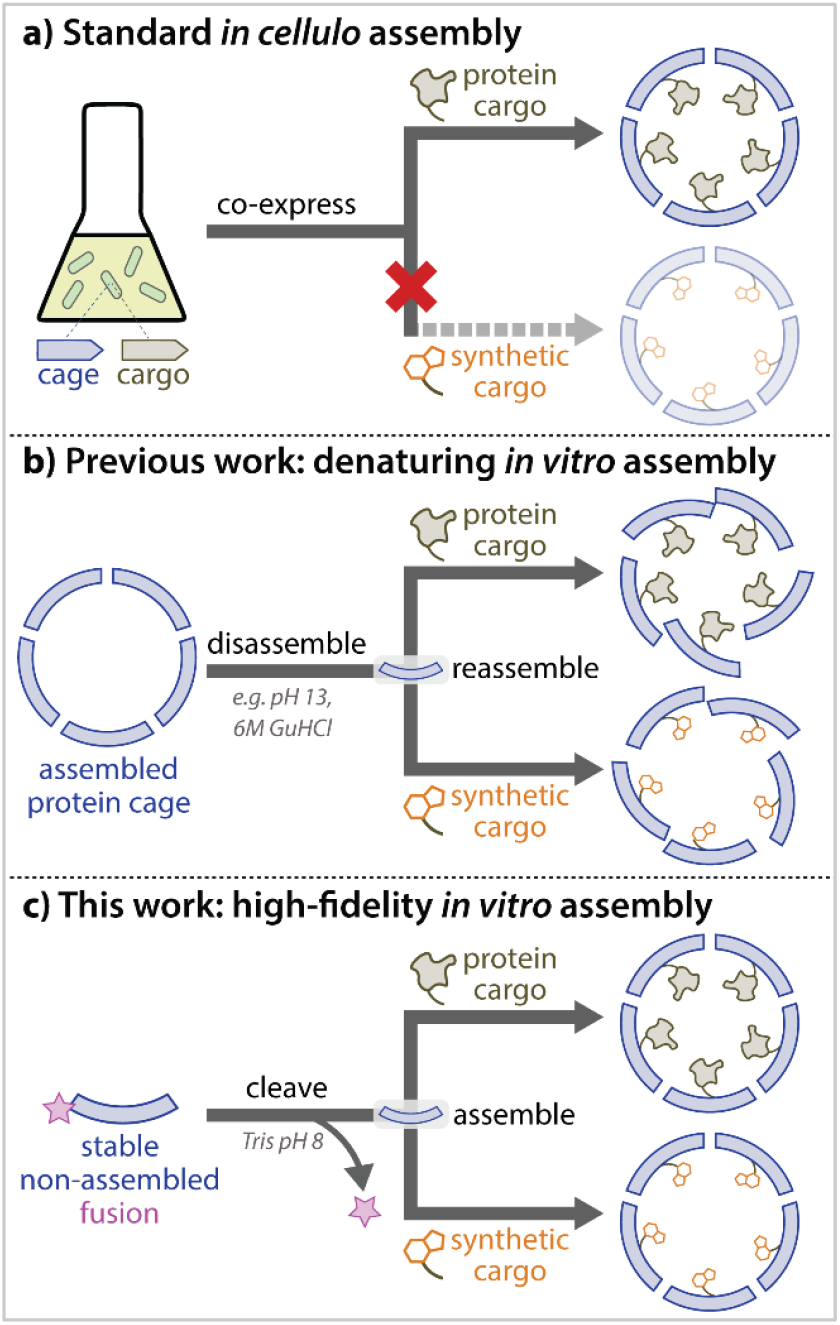
Strategies for packaging cargo into protein cages. **a)** Cargo packaging typically occurs in cells, where cargo that can be produced inside cells (proteins, nucleic acids) is packaged during the expression and assembly process. This method is generally incompatible with synthetic non-biological cargo. **b)** Current methods for *in vitro* packaging of synthetic cargo requires cages to be disassembled then subsequently reassembled in the presence of cargo, which can lead to significant assembly defects and aggregation. **c)** We report a fusion-based strategy for triggering *in vitro* assembly in encapsulin protein cages, producing highly uniform and thermostable cages that can be loaded with diverse synthetic cargo.

In this work, we report the high-fidelity *in vitro* packaging of diverse synthetic cargo into encapsulin protein cages. Encapsulins are simple non-viral protein cages with exceptional stability and a native peptide-mediated mechanism for loading proteinaceous cargo,^12^ serving as ideal candidates for engineering drug delivery vehicles,^41–43^ vaccine scaffolds,^41,44^ and nanoreactors.^14,45^ Until now, the ability to package synthetic cargo into encapsulin cages *in vitro* has been challenging, as fully-assembled encapsulins are not sufficiently porous to permit entry of larger molecular cargo.^16,46^ Previously reported protocols for *in vitro* packaging require exposure to extreme buffer conditions (e.g. pH 1, pH 13, or 7 M GuHCl) for cage disassembly,^47^ leading to low-yielding reassembly with substantial cage defects and aggregation (Figure 1b).^22,23^ Here, we achieve *in vitro* assembly and cargo packaging by using a protein fusion strategy to circumvent the need for disassembly (Figure 1c). This strategy outperforms current *in cellulo* assembly and *in vitro* disassembly-reassembly methods for encapsulins, producing cargo-filled cages with superior uniformity and thermostability.

## Results

### LanM-QtEnc is a stable fusion protein that does not readily self-assemble into native cages

To engineer a stable pre-assembly form of encapsulin, we fused the 12 kDa lanthanide-binding protein lanmodulin (LanM) to the internally facing *N*-terminus of the 32 kDa encapsulin protein from *Quasibacillus thermotolerans* (QtEnc). We chose QtEnc as it natively forms the largest known encapsulin protein cage, consisting of 240 protomers with an external diameter of 42 nm,48 while we chose LanM as the fusion partner due to its disordered structure (in the absence of lanthanides),49 potentially serving as a dynamic steric blockade to prevent the self-assembly of QtEnc. We fused the two proteins *via* a flexible glycine-serine linker and a TEV protease cleavage site, allowing for removal of the LanM fusion as a trigger for assembly. Finally, we included a hexahistidine tag at the *N*-terminus of LanM to facilitate purification. The resulting 46 kDa fusion protein is henceforth referred to as LanM-QtEnc.

Upon recombinant production in *E. coli*, we found that LanM-QtEnc is a stable fusion protein that does not self-assemble into cages. Purification of LanM-QtEnc involved an initial affinity chromatography step over Ni-NTA resin, followed by size-exclusion chromatography (SEC) on a Superdex 200 column to yield the pure fusion protein as determined by SDS-PAGE analysis (Figure 2a). Subsequent blue native PAGE (BN-PAGE) analysis showed that LanM-QtEnc predominantly forms a single low molecular weight species, in contrast to the expected 8 MDa cage from *in cellulo* assembly which is henceforth referred to as wild-type QtEnc (Figure 2b). Attempts to visualise the protein by negative-stain transmission electron microscopy (TEM) were consistent with lack of assembled cage-like structures (SI Figure S3.1a).

**Figure 2.**
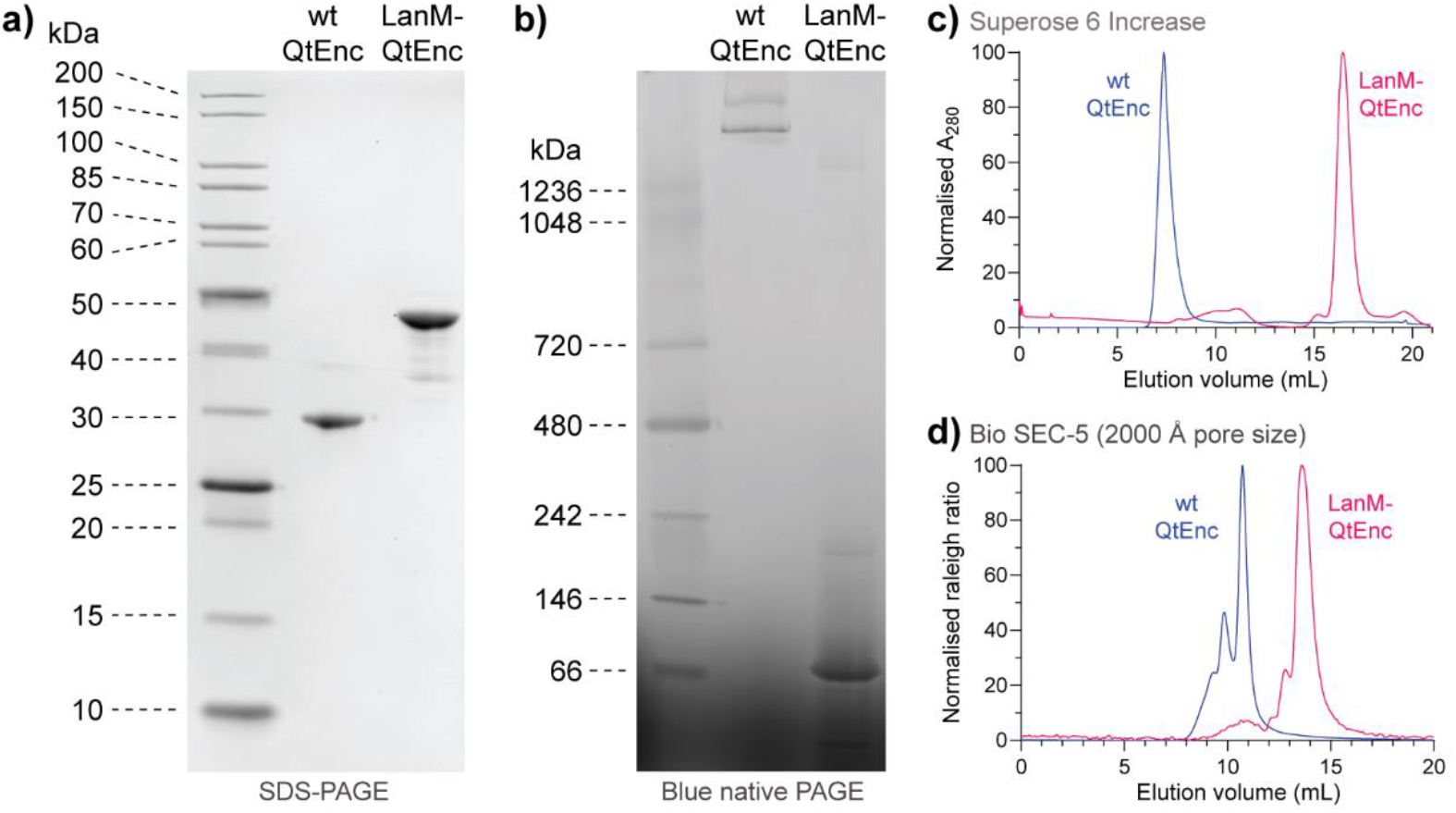
LanM-QtEnc is an encapsulin fusion that does not self-assemble into cages. **a)** SDS-PAGE analysis of purified wild-type QtEnc and LanM-QtEnc fusion shows the expected protein bands at 32 and 46 kDa respectively. **b)** Blue native PAGE analysis of LanM-QtEnc shows that the fusion exists predominantly in a low molecular weight form, while wild-type QtEnc forms the expected high molecular weight assembly, along with a faint smeared band corresponding to other potentially aberrant assemblies. **c)** Size-exclusion chromatography on a Superose 6 Increase 10/300 GL column confirms that LanM-QtEnc exists in an unassembled state, distinct from the cage assembly formed by wild-type QtEnc. **d)** Size-exclusion chromatography with light scattering detection on a Bio SEC-5 2000 Å HPLC column provides sufficient resolution and sensitivity at the megadalton size range to provide further evidence of aggregated or misassembled species in wild-type QtEnc.

We used analytical SEC to confirm that LanM-QtEnc exists in a predominantly unassembled form. SEC was performed on two different columns to achieve sufficient separation across a broad size range. Separation on a Superose 6 Increase column 10/300 GL (5-5000 kDa) showed a clear difference between wild-type QtEnc and LanM-QtEnc (Figure 2c), with the wild-type cage eluting early in the unretained peak, and the fusion eluting 9 minutes later, near the end of the chromatograph. Meanwhile, a Bio SEC-5 HPLC column with 2000 Å pore size (maximum mass range >10 MDa) also showed a substantial difference in retention time (Figure 2d), using light scattering in place of UV absorbance to enhance detection sensitivity at analytical scale. Notably for wild-type QtEnc, we also observed the presence of two additional peaks with earlier retention times (Figure 2d), indicating the existence of aggregated or misassembled cages that likely formed as a result of recombinant overexpression in *E. coli*. This polydisperse behaviour has not previously been reported for QtEnc in the literature due to limited resolving range of analytical SEC columns used in prior studies,48,50 although published DLS measurements suggest the existence of larger species (47.2 nm average diameter reported, 42 nm expected based on cryo-EM).51

### Cages assembled *in vitro* upon fusion cleavage are more uniform and stable than cages assembled *in cellulo*

We observed the high-fidelity *in vitro* assembly of encapsulin cages upon cleavage of the LanM-QtEnc fusion with TEV protease. SEC analysis on the Bio SEC-5 HPLC column revealed that the *in vitro* assembled cages were more uniform than wild-type QtEnc assembled in *E. coli*, forming a single species with a calculated mass of 8.2 ± 0.2 MDa from multi-angle light scattering (MALS) measurements (Figure 3a, SI Figure S3.1b). Encapsulin cages were isolated post-cleavage by SEC on a Superose 6 Increase 10/300 GL column, eluting in the unretained peak as expected for a large 8 MDa assembly (Figure 3b). BN-PAGE analysis provided further confirmation of the clean and complete conversion to an assembled state (Figure 3c), while SDS-PAGE analysis showed that cleavage had proceeded to >95% completion based on gel densitometry (Figure 3d).

**Figure 3.**
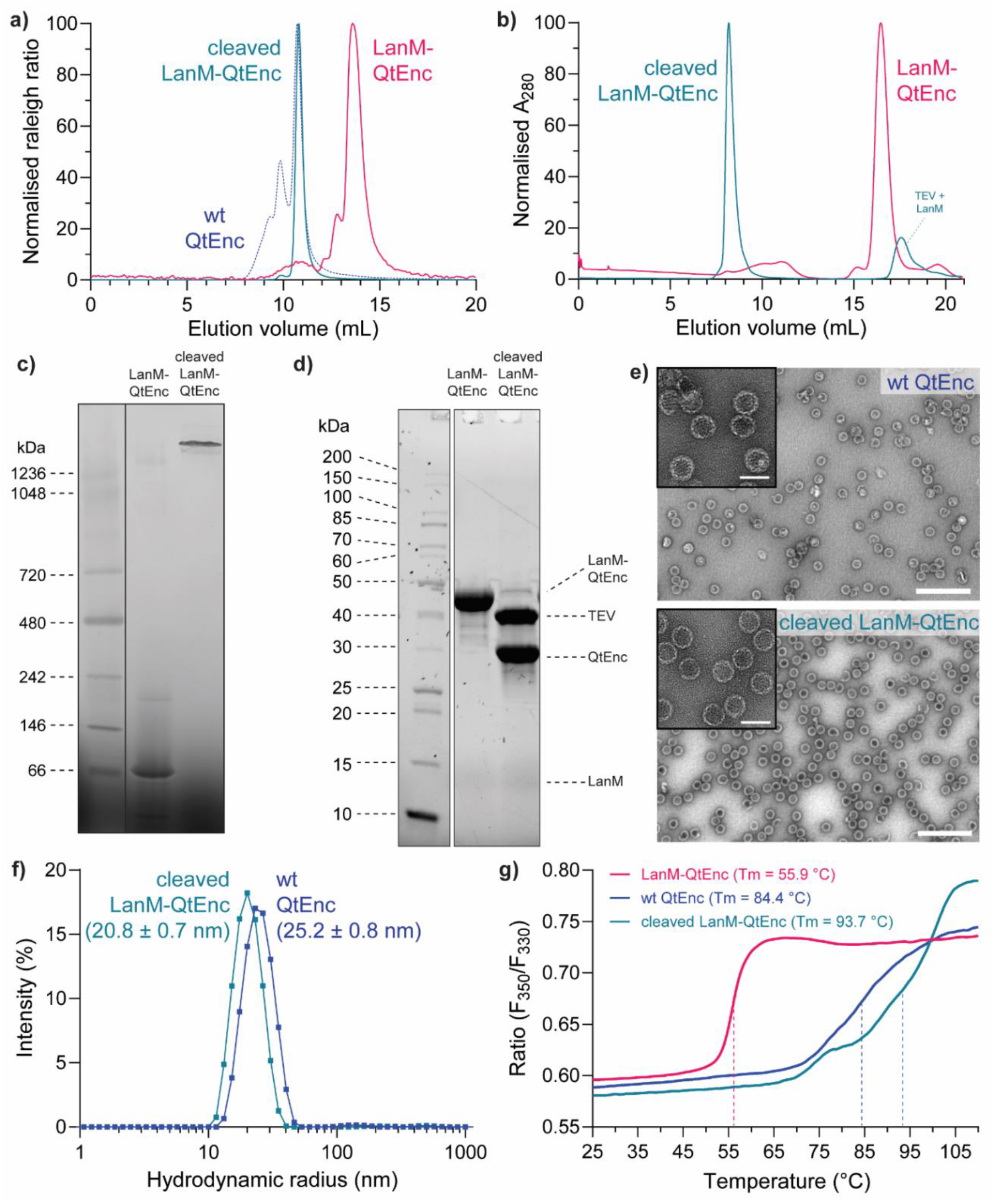
LanM-QtEnc cleavage triggers high-fidelity assembly of encapsulin cages. **a)** Analytical size-exclusion chromatography of the crude cleavage mixture on a Bio SEC-5 2000 Å HPLC column shows that the *in vitro* assembled encapsulins are highly monodisperse, without any of the additional peaks observed for wild-type QtEnc assembled in *E. coli*. **b)** Size-exclusion chromatography purification on a Superose 6 Increase 10/300 GL column shows clean conversion to a high molecular weight species. **c)** Blue native PAGE analysis of the purified *in vitro* assembled encapsulins indicates clean and quantitative conversion to a high molecular weight species. **d)** SDS-PAGE of the crude cleavage mixture shows that cleavage proceeded to >95% completion. **e)** Negative stain transmission electron microscopy of *in vitro* assembled encapsulins shows particles with similar cage morphology and size to wild-type QtEnc. The primary images were obtained at 40000× magnification (scale bar = 250 nm), while the inset images were obtained at 200000× magnification (scale bar = 50 nm). **f)** Dynamic light scattering measurements show that *in vitro* assembled encapsulins formed the expected size (42 nm diameter), while wild-type QtEnc was slightly larger due to the presence of minor aggregates or misassembled structures. **g)** Differential scanning fluorimetry shows that *in vitro* assembled encapsulins are more thermostable than wild-type QtEnc assembled in *E. coli*.

The successful assembly of uniform cages with the expected diameter of 42 nm was confirmed by negative stain TEM (Figure 3e) and calculated from hydrodynamic radius by dynamic light scattering (DLS) measurements (Figure 3f). The 50 nm diameter calculated for wild-type QtEnc by DLS provided further evidence of non-uniformity for assemblies produced in *E. coli*. Notably, unexpected diameters have been observed by DLS in previous attempts to disassemble and re-assemble encapsulins *in vitro* using denaturants, base, and heat,23 as well as *in vitro* studies on an engineered pH-switchable version of QtEnc,51 both of which were attributed to some degree of misassembly or aggregation.

The *in vitro* assembled cages showed exceptional thermal stability that exceeded that of the wild-type QtEnc assembled in *E. coli*. The melting temperature (T_m_) of the *in vitro* assemblies was 93.7 °C by differential scanning fluorimetry (DSF), compared to 55.9 °C for the LanM-QtEnc fusion prior to cleavage (Figure 3g). Meanwhile, wild-type QtEnc had a T_m_ of 84.4 °C, consistent with equivalent data from Giessen *et al*. who were the first researchers to characterise QtEnc (expressed and assembled in *E. coli*).48 This increased thermal stability is most likely a direct result of higher fidelity in the case of *in vitro* cage assembly.

### Protein cargo and synthetic molecules can be selectively and efficiently packaged during *in vitro* assembly

To demonstrate that the native cargo packaging system of encapsulins was functional *in vitro*, we encapsulated fluorescent protein mNeonGreen with the specific cargo loading peptide (CLP) for *Q. thermotolerans* encapsulin fused to the C-terminus (mNeon-Qt) (Figure 4a). The negative control for cargo packaging was the analogous mNeonGreen construct fused to the CLP for *T. maritima* encapsulin (mNeon-Tm), which does not mediate cargo packaging when co-expressed with QtEnc in *E. coli*.14 mNeon-Qt or mNeon-Tm was added during TEV cleavage of LanM-QtEnc, using a 1:10 molar ratio of cargo:encapsulin to avoid the possibility of sterically-induced defects arising from cargo overloading.52 After *in vitro* assembly, Ni-NTA resin was used to remove TEV protease, cleaved LanM, and any potentially unencapsulated mNeon cargo. SEC purification on a Superose 6 Increase 10/300 GL column showed that the efficiency of *in vitro* assembly was unaffected by the presence of cargo (Figure 4b), while negative stain TEM imaging confirmed the presence of uniform assemblies (Figure 4c). SDS-PAGE analysis showed that only mNeon-Qt co-eluted with the *in vitro* assembled encapsulins (Figure 4d), and BN-PAGE analysis confirmed that mNeon-Qt was selectively packaged into cages while mNeon-Tm was not packaged (Figure 4e). Unencapsulated mNeon-Qt was not observed during SEC purification (SI Figure S3.3), and absorbance measurements on purified mNeon-filled cages for total protein at 280 nm and mNeon-Qt cargo at 506 nm confirmed that mNeon-Qt had been packaged with minimal loss of cargo (SI Figure S3.4).

**Figure 4.**
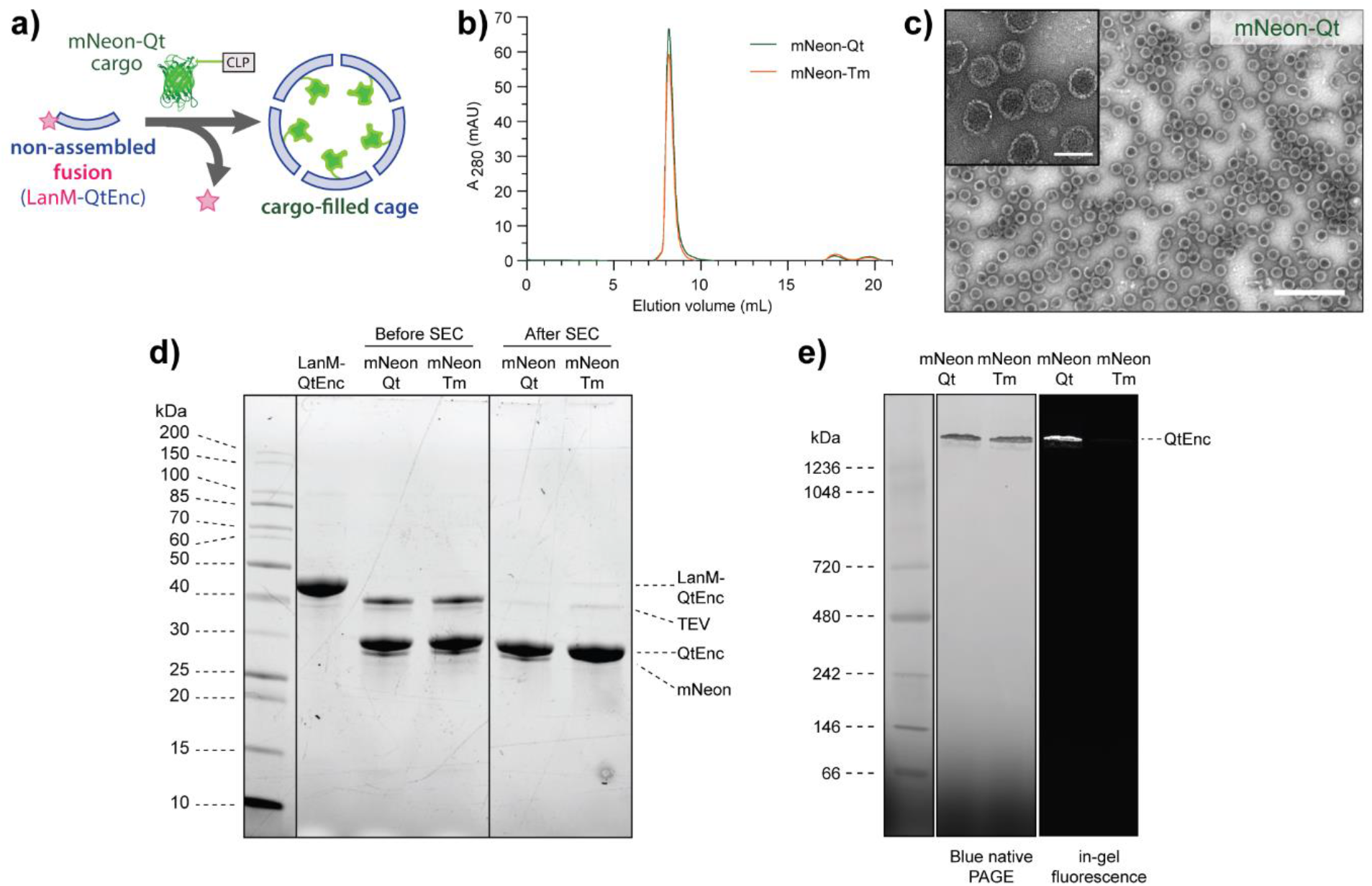
Selective packaging of protein cargo during *in vitro* assembly. **a)** Schematic of mNeon-Qt packaging into encapsulin by triggered *in vitro* assembly. **b)** Size-exclusion chromatography on a Superose 6 Increase 10/300 GL column shows that *in vitro* assembly is unaffected by the presence of cargo. **C)** Negative stain transmission electron microscopy of *in vitro* packaged encapsulin cages have similar morphology and size to wild-type QtEnc. The larger images were taken at 40000× magnification (scale bar = 250 nm), while the inset images were taken at 200000× magnification (scale bar = 50 nm). **d)** SDS-PAGE analysis of LanM-QtEnc cleavage in presence of cargo before and after SEC purification on a Superose 6 Increase 10/300 GL column. mNeon-Qt co-elutes with QtEnc cages while mismatched mNeon-Tm is not packaged. **e)** Blue native PAGE analysis shows clear in-gel fluorescence for *in vitro* assemblies containing mNeon-Qt protein cargo, while fluorescence is not visible for mNeon-Tm.

Next, we showed that synthetic small molecules can be packaged *in vitro* (Figure 5a). The synthetic fluorescent dye 5-carboxytetramethylrhodamine (TMR) was attached to the *N*-terminus of the CLPs for *Q. thermotolerans* (TMR-Qt) and *T. maritima* (TMR-Tm) by Fmoc solid-phase peptide synthesis. The *in vitro* cargo loading experiments were repeated with TMR in place of mNeonGreen and using Capto Core 700 resin in place of Ni-NTA resin after assembly to remove TEV protease, cleaved LanM, and any potentially unencapsulated cargo. Once again, the presence of synthetic cargo did not affect the fidelity and efficiency of assembly according to analytical SEC (Figure 5b) and TEM (SI Figure S3.5). In-gel fluorescence was observed after conducting BN-PAGE on purified cages *in vitro* loaded with TMR-Qt, confirming successful and specific encapsulation of the synthetic dye, while no fluorescence was observed for the TMR-Tm negative control (Figure 5c, SI Figure S3.6).

**Figure 5.**
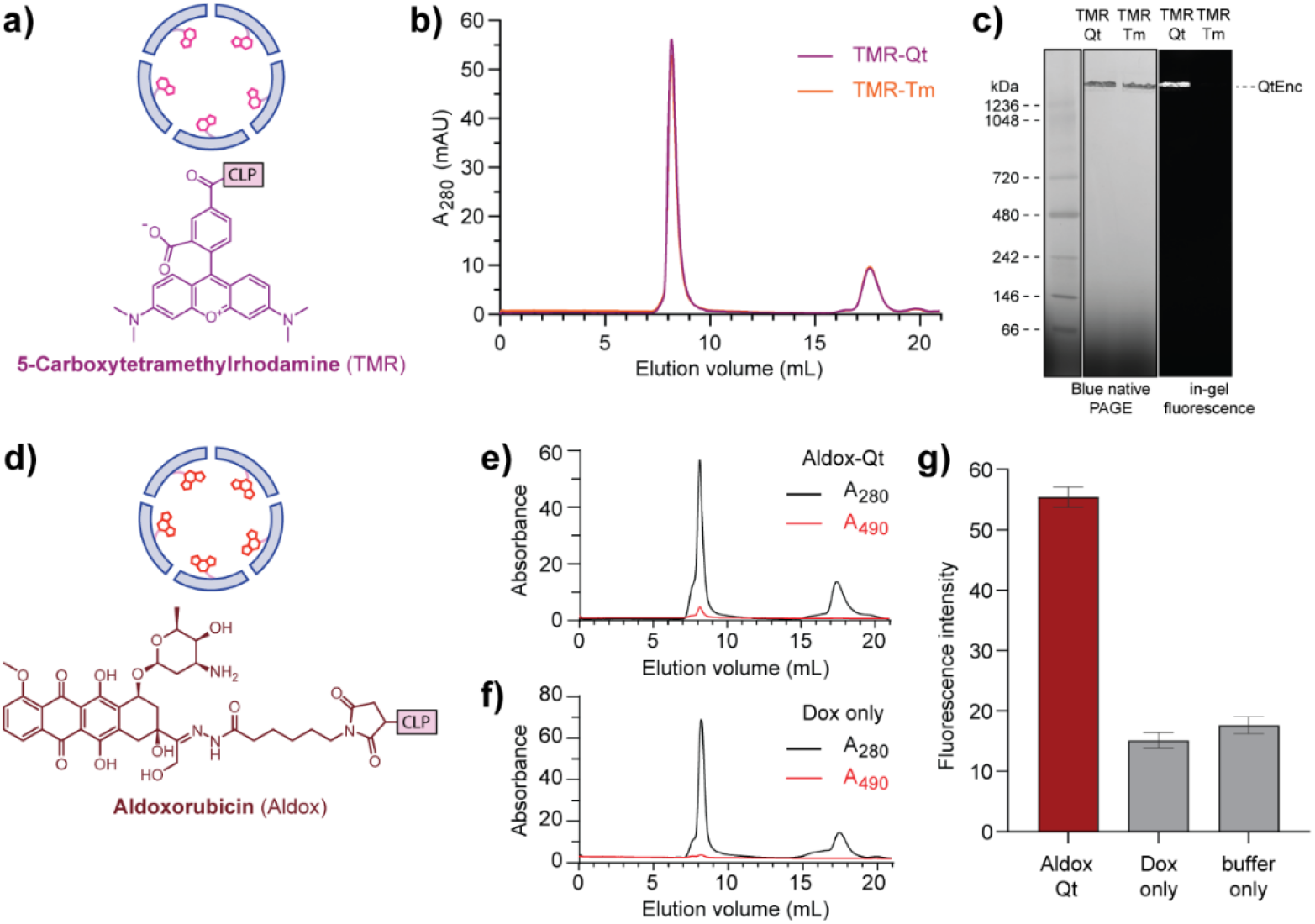
*In vitro* packaging of diverse synthetic cargo into encapsulin cages. **a)** Schematic of TMR-Qt *in vitro* packaged into encapsulin cages. **b)** Size-exclusion chromatography monitoring absorbance at 280 nm on a Superose 6 Increase 10/300 GL column shows that the fidelity of *in vitro* assembly is unaffected by the presence of TMR cargo. **c)** Blue native PAGE analysis shows clear in-gel fluorescence for *in vitro* assemblies with TMR-Qt synthetic cargo selectively packaged, while fluorescence is not visible for TMR-Tm. **d)** Schematic of Aldox-Qt *in vitro* packaged into encapsulin cages. **e-f)** Size-exclusion chromatography monitoring absorbance at 490 nm on a Superose 6 Increase 10/300 GL column shows that Aldox-Qt co-elutes with encapsulin assemblies, while doxorubicin without the CLP does not co-elute with encapsulin. **g)** Fluorescence intensity of the encapsulin fractions from SEC purification show aldoxorubicin fluorescence (emission at 595 nm) when Aldox-Qt is packaged, while the negative control of packaging doxorubicin without the CLP only shows background levels of fluorescence. Error bars correspond to the standard error of the mean across technical measurement replicates.

To expand the scope of cargo packaging beyond proteins and synthetic dyes, we packaged the synthetic drug molecule aldoxorubicin (Aldox), a maleimide-functionalised prodrug form of the chemotherapeutic doxorubicin (Dox) (Figure 5d). Aldox was conjugated onto a Cys-modified Qt CLP to generate Aldox-Qt, while free Dox was used as a negative control. Both Aldox-Qt and free Dox were *in vitro* packaged in a 2:1 molar ratio of encapsulin to cargo. This higher ratio of cargo packaging was used to provide sufficient photophysical signal for detection, given the reduced brightness of Aldox relative to the previously packaged fluorophores. Analytical SEC on a Superose 6 Increase 10/300 GL column showed an Aldox-specific absorbance peak (490 nm) in the encapsulin fraction for *in vitro* packaged Aldox-Qt (Figure 5e), while no Aldox absorbance was observed for *in vitro* packaged free Dox (Figure 5f). Furthermore, fluorescence emission was only observed in encapsulin samples purified after *in vitro* assembly with Aldox-Qt, while the negative control with free Dox showed equivalent signal to the buffer-only control (Figure 5g), further verifying that Aldox-Qt was selectively packaged. Taken together, these results show that diverse synthetic cargo can be efficiently packaged into encapsulins *in vitro* by taking advantage of the highly specific encapsulin-CLP interaction.

### Synthetic cargo packaged within encapsulins can be delivered into murine RAW 264.7 cells

We investigated whether *in vitro* assembled encapsulins could facilitate uptake of fluorescent mNeon-Qt and TMR-Qt cargo into live cells. Cellular uptake of packaged cargo into murine RAW 264.7 macrophages was assessed by confocal fluorescence microscopy. Intracellular fluorescence was readily observed in 83% of cells when encapsulated mNeon-Qt was added, while no fluorescence was observed when free non-encapsulated mNeon-Qt was used (Figure 6a-c). Similarly, intracellular fluorescence was evident for packaged TMR-Qt in 50% of cells, while free TMR-Qt did not result in any fluorescence (Figure 6d-f). This result suggests that encapsulation either enhances cellular uptake of fluorescent cargo or protects the cargo against degradation by the cellular machinery prior to imaging. In summary, these results provide a proof of concept that *in vitro* assembled encapsulins are viable cargo delivery vehicles.

**Figure 6.**
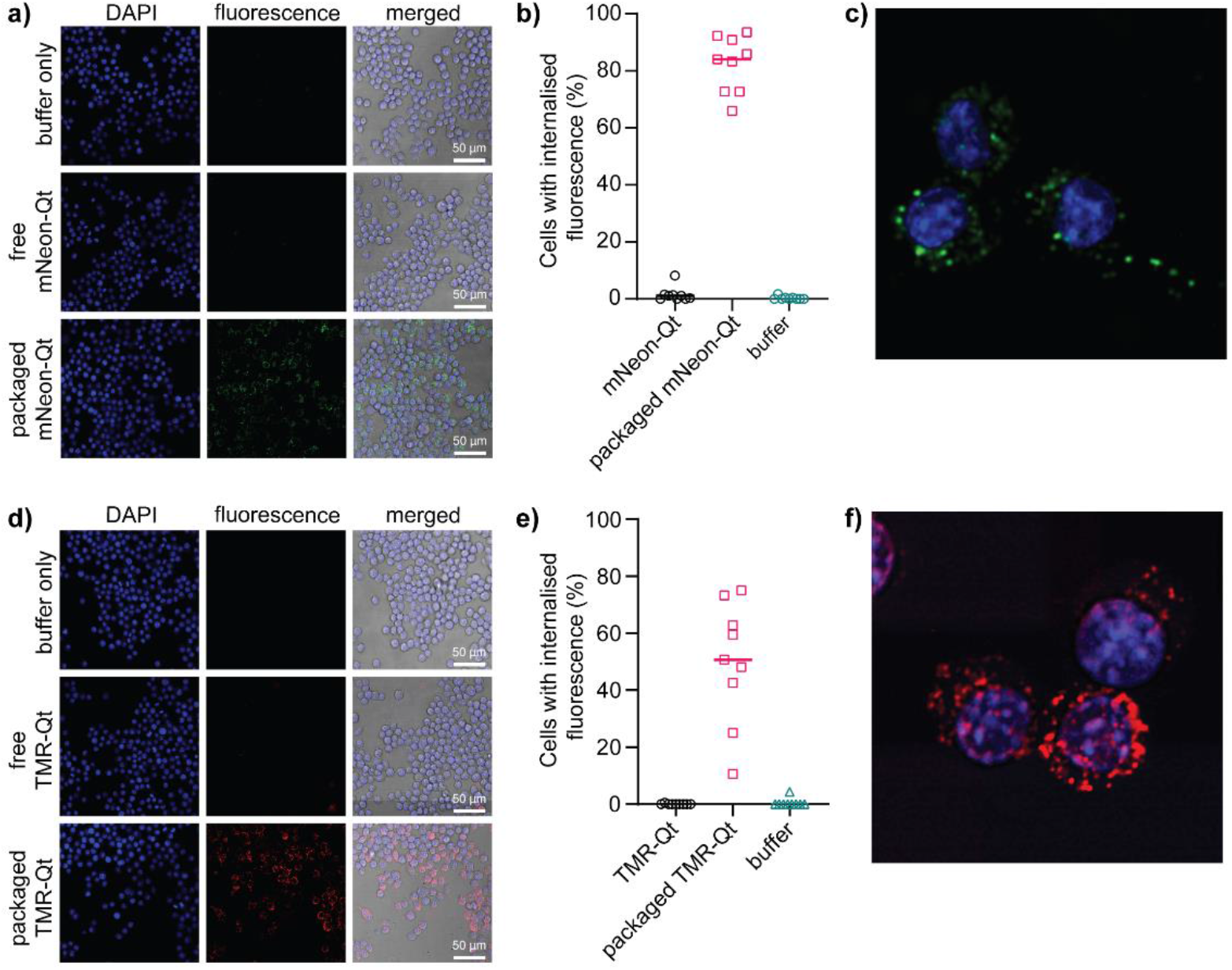
Cellular uptake of cargo *in vitro* packaged inside encapsulins. **a)** Confocal fluorescence microscopy images of murine RAW 264.7 cells with DAPI fluorescence (nuclear stain) and fluorescence in the FITC channel (mNeon). Merged indicates an overlay of the two channels, as well as brightfield imaging. *In vitro* packaged mNeon-Qt was observed inside cells while free mNeon-Qt was not observed. **b)** ImageJ quantification of fluorescence shows that a median of 83% of cells showed uptake of packaged mNeon-Qt, while minimal uptake was observed for free mNeon-Qt and buffer-only controls. Data analysis was performed on three independent fields of view from three biological replicates. **c)** Representative magnified image of RAW 264.7 cells showing intracellular localisation of packaged mNeon-Qt. **d)** Confocal microscopy images of RAW 264.7 cells with DAPI and fluorescence in the TRITC channel (TMR). *In vitro* packaged TMR-Qt was observed inside cells while free TMR-Qt was not observed. **e)** ImageJ quantification of fluorescence shows that a median of 50% of cells showed uptake of packaged TMR-Qt, while minimal uptake was observed for free TMR-Qt and buffer-only controls. Data analysis was performed on three independent fields of view from three biological replicates. **f)** Representative magnified image of RAW 264.7 cells showing intracellular localisation of packaged TMR-Qt.

## Discussion

LanM-QtEnc is a novel protein construct that enables *in vitro* packaging of synthetic cargo into encapsulin cages, providing access to new biotechnological opportunities that cannot be explored using standard cell-based assembly approaches. *In vitro* packaging is necessary for any synthetic molecular cargo that is incompatible with living cells due to cytotoxicity, metabolic instability, or membrane impermeability. In applications where multiple cargo types are packaged simultaneously, *in vitro* packaging also provides the opportunity for far greater control over cargo stoichiometry than state-of-the-art genetic methods for tuning expression levels in cells.^53,54^ Even in the case of biomolecular cargo that can be produced in cells, *in vitro* packaging solves any timing issues where cargo requires additional processing prior to packaging, such as any cargo that requires multiple subunits to form an active complex, or cargo that must undergo post-translational modification to be functional.

A distinctive feature of the LanM-QtEnc system is the exceptional fidelity of cage assembly and cargo packaging. Defective assembly pathways are commonly encountered during most *in vitro* assembly methods,^23,26,55–57^ as well as in some *in cellulo* assembly methods.^58^ Low-fidelity assembly processes result in defects or aggregates that can reduce the overall yield of desired cages, pose challenges for sample purification, and undermine the fundamental properties that make protein cages valuable in biotechnology. In nanoreactor design for example, protein cages can act as a semi-permeable barrier to control substrate flux and sequester toxic or unstable intermediates, or alternatively, act as a protective layer for fragile enzymatic cargo.^14,59–64^ Assembly defects compromise the ability of cages to achieve both of these core functions. In drug delivery applications, heterogeneous formulations arising from defective cage assemblies may be less stable *in vivo*, and hence less effective in shielding cytotoxic payloads from non-specific release prior to reaching the site of action. In addition, cargo packaging efficiency is a critical parameter when working with valuable cargo where losses must be minimised.

The LanM-QtEnc fusion stands out as a unique example of a stable encapsulin construct that does not form a fully-assembled cage. In most literature examples of encapsulin engineering, the assembled state of encapsulins is strongly favoured. Lee *et al*. have reported several *N*-terminal HBCM2 fusion constructs to engineered versions of the 24 nm encapsulin from *T. maritima*, observing cage formation along with increased heterogeneity in most cases.^65^ A notable native example of *N*-terminal encapsulin fusion that does not impede assembly is the encapsulin system from *Pyrococcus furiosus*, which has its cargo directly fused on the *N*-terminal interior of the cage.^66^ The *N*-terminal fusion behaviour of LanM-QtEnc also differs from the HK97 bacteriophage capsid protein (from which encapsulins derive their HK97-like protein fold), where the scaffold protein gp5 is initially fused to the capsid protein during the immature Prohead-I that can form a mix of cages and dissociated capsomeres, but then is cleaved to initiate maturation to the final Head II phage capsid form.^67^

While the robustness of assembled encapsulin cages is ideal for many biotechnological applications, this robustness has also posed a significant technical barrier for previously reported disassembly-reassembly protocols, necessitating extreme pH or chemical denaturants to attempt cargo loading with poor fidelity.^22,43^ In one exception, Jessen-Trefzer and co-workers covalently conjugated synthetic small molecule cargo on the inside of encapsulin cages without any disassembly,^68^ although larger cargo is expected to be excluded from accessing the cage interior due to the narrow pore size of most encapsulins. Meanwhile, Giessen and co-workers inserted a pH-sensitive GALA peptide into the encapsulin sequence to generate a dynamic switchable assembly, albeit with an unexpected increase in diameter suggesting possible heterogeneity (62.2-76.8 nm average diameter by DLS).^51^ Unlike all these examples, LanM-QtEnc entirely circumvents the challenge by starting from a non-assembled state. More generally, the extreme buffer conditions required for encapsulin disassembly are not shared by all protein cages. Some VLPs are inherently capable of disassembly *via* relatively mild changes in experimental buffer conditions. VLPs of the brome mosaic virus coat protein can assemble at low pH and ionic strength but disassemble at neutral pH and high ionic strength,^69^ while VLPs of polyomaviruses capsid proteins such as SV40 VP1 can be isolated from cells as pentameric capsomeres and assembled with the addition of DNA.^70^ As a trade-off however, these VLPs have inherently less tolerance to the wide range of non-native conditions required for different biotechnological applications.

Beyond encapsulins and VLPs, there has been recent surging interest in the *in vitro* assembly of other types of protein cages. In work on the *Haliangium ochraceum* microcompartment shell protein, Kerfeld and co-workers used a SUMO-fusion strategy for the *in vitro* assembly of tubes and cages, where SUMO fusion produced a soluble hexameric state and subsequent SUMO cleavage led to higher-order oligomeric assembly.^71^ Hilvert and co-workers recently used a fusion of maltose-binding protein to achieve *in vitro* assembly of an engineered lumazine synthase cage that packages RNA,^72^ while Podobnik and co-workers used this strategy to assemble hollow nanotubes based on the coat protein of potato virus Y.^73^

In addition to building nanoreactors and drug delivery systems based on *in vitro* assembly, there are multiple future avenues for deepening our fundamental understanding of the LanM-QtEnc fusion system. In future work, we will investigate the origins of LanM stabilisation of the non-assembled state, to determine whether LanM is indeed unique in this role and whether disorder of the fusion partner is a significant contributor. The impact of lanthanide binding to LanM on self-assembly and cage stability is another possible area of further investigation. Finally, we will investigate whether LanM fusion can be used on other encapsulins, including pore-engineered encapsulins, and potentially even more generally across other HK97-fold capsids and unrelated cages, ultimately leading to a diverse set of *in vitro* assembled protein cages for new biotechnological applications.

## Methods

### Molecular cloning

The codon-optimised gene encoding LanM-QtEnc was synthesised as a gBlock (Integrated DNA Technologies) and cloned into a pETDuet-1 vector between the NcoI and KpnI restriction sites by Gibson assembly using NEBuilder HiFi DNA Assembly Master Mix (New England Biolabs). Constructs were transformed into DH5α *E. coli* cells and the gene sequence was verified by Sanger sequencing at the Australian Genome Research Facility. Full details of DNA sequences are provided in the Supporting Information (SI Section 1).

### Recombinant protein production

Plasmids were transformed into BL21(DE3) *E. coli*. Single colonies were grown overnight at 37 °C in lysogeny broth (LB) as 10 mL cultures. Overnight cultures were used to inoculate 400 mL LB cultures (OD600 0.05) and grown at 37 °C to OD600 0.4-0.6, followed by induction with 0.1 mM isopropyl β-D-1-thiogalactopyranoside (IPTG) and expression overnight at 18 °C. Cells were pelleted at 3900 *g* for 20 min (5810R centrifuge with S-4-104 rotor, Eppendorf) and stored at -20 °C until purification.

### Protein purification

The following method was used for the purification of LanM-QtEnc, mNeon-Qt, and mNeon-Tm. Frozen pellets were resuspended in 25 mL SEC buffer (50 mM Tris pH 8, 200 mM NaCl, 1 mM tris(2-carboxyethyl)phosphine (TCEP)) with the addition of 10 µg/mL DNase I (bovine pancrease, Sigma-Aldrich), 100 µg/mL lysozyme (chicken egg white, Sigma-Aldrich), and 1× protease inhibitor (cOmplete EDTA-free Protease Inhibitor Cocktail, Roche). After 30 min on ice, cells were lysed by sonication using a Sonopuls HD 4050 with TS-106 probe (Bandelin) with a program of 55% amplitude for 11 min with a pulse time of 8 s on and 10 s off. The lysate was clarified at 17000 *g* for 40 min, then incubated with rocking for 1 h at 4 °C with 1 mL HisPur Ni-NTA resin (ThermoFisher Scientific) pre-equilibrated in SEC buffer. The mixture added to an Econo-Column chromatography column (Bio-Rad), and the beads were washed with 3 × 2 mL wash buffer (50 mM Tris pH 8, 200 mM NaCl, 1 mM TCEP, 20 mM imidazole), and eluted with 4 × 1 mL elution buffer (50 mM Tris pH 8, 200 mM NaCl, 1 mM TCEP, 500 mM imidazole). The eluted protein was purified by size-exclusion chromatography on a HiLoad 16/600 Superdex 200 pg column using SEC buffer at a flow rate of 1 mL/min on an AKTA Pure (Cytiva). All fractions containing the desired protein were concentrated and buffer exchanged using Amicon Ultra-15 centrifugal filters (10 kDa MWCO, Merck) into cleavage buffer (50 mM Tris pH 8, 50 mM NaCl, 1 mM TCEP, 0.5 mM ethylenediaminetetraacetic acid (EDTA)).

Tobacco Etch Virus (TEV) protease was purified using the same lysis and Ni-NTA affinity purification protocol. The eluted protein was dialysed into TEV-SEC buffer (20 mM HEPES pH 8, 200 mM NaCl, 2 mM EDTA, 2 mM dithiothreitol (DTT)) using Snakeskin dialysis tubing (3.5 kDa MWCO, ThermoFisher Scientific). The dialysed protein was purified by size-exclusion chromatography on a HiLoad 16/600 Superdex 200 pg column using TEV-SEC buffer at a flow rate of 1 mL/min on an AKTA Pure (Cytiva). Fractions containing TEV protease were concentrated and buffer exchanged using Amicon Ultra-15 centrifugal filters (10 kDa MWCO, Merck) into 40 mM HEPES pH 8, 400 mM NaCl, 4 mM EDTA, 4 mM DTT, and diluted 1:1 with glycerol before being stored at -80 °C.

Wild-type QtEnc was purified using the same lysis conditions (while omitting TCEP from the lysis buffer), but following lysate clarification, solid ammonium sulfate was added to the supernatant to 20% (w/v) saturation. The mixture was rocked at 4 °C for 15 minutes prior to centrifugation at 10,000 x g for 15 minutes. Following clarification, solid ammonium sulfate was added to a total of 50% (w/v) saturation. This mixture was rocked at 4 °C for 15 minutes followed by another centrifugation at 10,000 x g for 15 minutes. The pellet was isolated and resuspended in 5 mL of purification buffer (50 mM Tris pH 8, 200 mM NaCl). This sample was applied to Snakeskin Dialysis Tubing (3.5 kDa MWCO, ThermoFisher Scientific) and dialysed against 1 L of purification buffer at 4 °C for 2 – 4 h followed by overnight dialysis against a fresh 1 L of purification buffer under the same conditions. The following morning, the sample was recovered and nucleic acids were removed via anion-exchange chromatography on a HiPrep Q XL 16/10 column at a flow rate of 5 mL/min using an AKTA Start (Cytiva). The unretained peak was collected. Following this, the sample was purified through two rounds of size-exclusion chromatography using an AKTA Pure (Cytiva). The first step used a HiPrep 16/60 Sephacryl S-500 HR column and the QtEnc-containing fractions were concentrated using Amicon Ultra-15 filters (100 kDa MWCO, Merck) before being applied to a Superose 6 Increase 10/300 GL column. The encapsulin-containing fractions were quantified via absorbance at 280 nm and stored at 4 °C until further use.

### Solid-phase peptide synthesis

Manual peptide synthesis was performed on Rink Amide MBHA resin (0.52-0.65 mmol/g loading, Merck). Couplings were carried out by adding HATU (4 eq) and DIPEA (4 eq) to a solution of the Fmoc-protected amino acid (4 eq) in DMF. This pre-activated mixture was then added to the resin in DMF and shaken for 1 h. The side chain protecting groups used were: t-Bu for Asp, Glu, Ser, Thr; Boc for Lys; Pbf for Arg; Trt for Asn, Gln, His. Fmoc deprotection was carried out with 20% piperidine in DMF (2 × 1 min, 1 × 10 min). N-terminal capping with 5-TAMRA was conducted by adding pre-activated 5-TAMRA (4 eq) with HATU (4 eq) and DIPEA (8 eq) in DMF to the resin swelled in DMF and shaking at rt for 3 h.

Cleavage from the resin was achieved with TFA containing 2.5% triisopropylsilane and 2.5% water for 2 h. After cleavage, the mixture was concentrated under a stream of nitrogen. The crude residue was triturated with diethyl ether (15-20 mL) before purification by reverse-phase chromatography.

Semi-preparative reverse-phase HPLC was performed on a Waters 1525 binary HPLC pump with a Waters 2707 autosampler. Peptide samples were eluted through a Waters XBridge Peptide BEH C18 OBD™ Prep Column (10 mm × 250 mm, 5 μm) using a linear gradient system of 0.1% (v/v) TFA in MilliQ water (solvent A) and 0.1% (v/v) TFA in MeCN (solvent B) at a flow rate of 4 mL/min over 30 min. Peptides were monitored by UV absorbance at 220 nm on a Waters 2489 UV Detector with semi-prep cell. Fractions were collected with Waters Fraction Collector III.

To produce Aldox-Qt, a Cys-modified Qt CLP was synthesised and purified as per the aforementioned methodology, with the addition of an *N*-terminal cysteine residue. The peptide was then capped by adding acetic anhydride (4 eq) and DIPEA (4 eq) in DMF for 1 h before cleavage from the resin and HPLC purification as before. The purified peptide in DMSO (1 eq), aldoxorubicin in DMF (1 eq), and DIPEA (1 eq) were shaken in 1:1 MeCN:water at 37 °C for 3 h to yield crude Aldox-Qt. Automated flash reverse-phase column chromatography was carried out on a Biotage Selekt using pre-packed Biotage Sfär C4 column (10 g) for the purification of Aldox-Qt.

LCMS chromatograms were obtained using a Shimadzu Nexera-I LC-2040C Plus coupled to a Shimadzu LCMS-2020 mass spectrometer ESI single quadrupole mass detector. Peptide samples were eluted through a Shimadzu Shim-Pack Velox SP-C18 column (2.1 mm × 50 mm, 2.7 µm) with a linear gradient system of 0.1% (v/v) formic acid in MilliQ water (solvent A) and 0.1% (v/v) formic acid in MeCN (solvent B) at a flow rate of 0.4 mL/min over 6.4 min. Peptide samples were monitored by UV absorbance at 220 and 254 nm. Mass spectra were obtained by electrospray ionisation in both positive and negative modes, scanning between m/z 200 and 2000.

### In vitro encapsulin assembly and cargo loading

Cleavage reactions were performed on 100 µL scale by incubating 2 mg/mL LanM-QtEnc and 0.4 mg/mL TEV protease in cleavage buffer overnight at 34 °C. Cargo loading experiments included a mixture of LanM-QtEnc and either mNeon or TMR cargo in a 10:1 molar ratio, or Aldox cargo in a 2:1 molar ratio. After overnight cleavage, the reactions were passed through 0.1 mL Ni-NTA resin (for empty or mNeon loading), Capto Core 700 resin (for TMR loading). The *in vitro* assemblies were then purified by size-exclusion chromatography on a Superose 6 Increase 10/300 GL column at 0.5 mL/min flow rate in SEC buffer.

The packaging efficiency of mNeonGreen was estimated by measuring the absorbance of purified cargo-filled encapsulins at 506 nm on a Nanodrop ND-1000 spectrophotometer (ThermoFisher Scientific). The extinction coefficient of mNeonGreen at 506 nm (116,000 M-1 cm-1) was used to calculate the concentration in packaged samples.74 The packaging of Aldox was monitored by absorbance at 490 nm during the SEC purification. Fluorescence of the SEC purified Aldox loaded shells was measured using a Perkin Elmer EnSpire multimode plate reader (excitation 470 nm, emission 595 nm). Due to low signal intensity, 100 technical replicate measurements were obtained for each sample, with the mean intensity reported with the standard error.

### Analytical size exclusion chromatography

Analytical SEC was conducted on a Nexera Bio HPLC (Shimadzu) using a Bio SEC-5 2000 Å pore size column (7.8 × 300 mm, Agilent) with a Bio SEC-5 2000 Å guard column (7.8 × 50 mm, Agilent). Multi-angle light scattering (MALS) measurements were obtained on a Dawn 8 MALS detector (Wyatt Technologies). Concentration was calculated from change in refractive index assuming a dn/dc of 0.185. The mobile phase was phosphate-buffered saline (137 mM NaCl, 2.7 mM KCl, 10 mM Na2HPO4, 1.8 mM KH2PO4, pH 7.4) at a flow rate of 1 mL/min for 20 min, analysing 20 µL samples taken directly from cleavage reactions. Astra software v7.3.0.18 (Wyatt Technologies) was used to calculate molecular weights.

### Polyacrylamide gel electrophoresis

SDS-PAGE was conducted using Mini-PROTEAN TGX Stain-Free gels (Bio-Rad) with Tris-glycine-SDS running buffer. Gels were stained using Coomassie Brilliant Blue G-250 (80 mg/L in 30 mM HCl). Unstained Protein Standard, Broad Range 10-200 kDa (New England Biolabs) was used as the ladder for all SDS-PAGE gels.

Blue Native PAGE was conducted using NativePAGE 3-12% Bis-Tris 1.0 mm Mini Protein Gels (ThermoFisher Scientific). Protein samples were diluted to an A280 of 0.4 in NativePAGE sample buffer and 9 µL was loaded in each well. NativeMark Unstained Protein Standard (ThermoFisher Scientific) was used as the ladder for all BN-PAGE gels. The anode, dark blue cathode, and light blue cathode buffers were prepared as per manufacturer instructions. Gels were run in dark blue buffer for 5 minutes at 150 V before switching to light blue buffer for 75 minutes at 150 V. The anode buffer was not changed during this time. After electrophoresis, in-gel fluorescence was measured for samples loaded with TAMRA (Trans-UV excitation 302 nm, 590/110 nm emission filter) or mNeonGreen (Epi-blue excitation 460-490 nm, 532/28 nm emission filter) on a ChemiDoc MP Imaging System (Bio-Rad). Gels were then stained with Coomassie Brilliant Blue G and then destained in water prior to imaging.

### Transmission electron microscopy

Encapsulin samples for negative-stain TEM were diluted to 100 μg/ml in 20 mM Tris buffer (pH 8). Gold grids (200-mesh coated with Formvar-carbon film, EMS) were made hydrophilic by glow discharge at 50 mA for 30 s. The grid was floated on 20 μl of sample for 1 min, blotted with filter paper, and washed once with distilled water before staining with 20 μl of uranyl acetate for 1 min. TEM images were captured using a JEOL 1400 microscope at 120 keV at Sydney Microscopy and Microanalysis.

### Dynamic light scattering and thermal unfolding analysis

Wt QtEnc and *in vitro* assembled QtEnc were prepared at 0.5 mg/mL in 50 mM Tris pH 8, 200 mM NaCl and loaded onto Prometheus Panta (NanoTemper) as per the manufacturer instructions. Samples were analysed using the Size Analysis function, in which isothermal DLS scans (10 acquisitions, 5s each, 100% DLS-Laser intensity) were performed at 25 °C.

For differential scanning fluorimetry (nanoDSF), wt QtEnc, LanM-QtEnc and *in vitro* assembled QtEnc were prepared at 0.5 mg/mL in the same buffer and loaded onto standard capillaries (NanoTemper). Capillaries were sealed at the ends (Capillary Sealing Paste; NanoTemper). Thermal unfolding was determined on Prometheus Panta (NanoTemper) with heating ramp of 1 °C from 25 °C to 110 °C and 20% sensitivity setting.

### Cell culture and in vitro loaded QtEnc uptake into murine cells

Murine RAW 264.7 cells were maintained in DMEM (Gibco) supplemented with 10% fetal bovine serum (FBS) and 1% penicillin/streptomycin at 37 °C with 5% CO2.

Cells were seeded (day 1) at a density of 50,000 cells/well in black 96 well plates (ThermoFisher Scientific) overnight. On day 2, culture media was removed, cells washed with PBS, and media was replaced with starvation media (culture medium with 0% FBS). On day 3, starvation media was removed, cells were washed, and normal culture media containing 0.4 µg/ml of cargo-filled QtEnc, or equivalent control, was added to the cells. Cells were incubated with cargo-filled QtEnc for 2 h at 37 °C with 5% CO2. Following incubation, cells were washed three times with PBS with agitation for 5 mins at room temperature. Cells were fixed by incubating the cells with 4% paraformaldehyde for 15 mins with agitation at room temperature. Cells were then washed again and counterstained with DAPI. Following a third wash, 100 µL of PBS was then added to the cells in preparation for imaging. The proportion of cells with internalised fluorescent cargo-filled QtEnc was visualized *via* confocal microscopy using an A1 confocal microscope (Nikon). Three independent fields of view were obtained for each of three biological replicates, and the resulting images were analysed in Fiji (ImageJ).

## Acknowledgements

YHL acknowledges funding from the Australian Research Council (DE19010062, DP230101045) and Westpac Scholars Trust (WRF2020). We acknowledge the core facilities at Sydney Analytical and Sydney Microscopy and Microanalysis for providing infrastructure support. TNS, RS, LSRA, and YHL acknowledge support from the ARC Centre of Excellence in Synthetic Biology, while YHL also acknowledges support from the ARC Centre of Excellence for Innovations in Peptide and Protein Science. AC acknowledges funding from the Dementia Australia Research Foundation; Mason Foundation; and the National Foundation for Medical Research and Innovation. The authors acknowledge the use of the Nikon A1 inverted confocal microscope in the Microbial Imaging Facility at AIMI in the Faculty of Science, the University of Technology Sydney.

