## Supplementary Information for "High-fidelity *in vitro* packaging of diverse synthetic cargo into encapsulin protein cages"

**Contents**

1. Gene fragment sequences
2. Protein sequences
3. Supplementary SEC data, PAGE gels and TEM images
4. Peptide synthesis and characterisation

### 1. Gene fragment sequences

#### 1.1 Codon-optimised sequence of wild-type QtEnc gene

The synthetic gene was inserted into a pETDuet-1 vector between NcoI and BamHI restriction sites by Gibson assembly.

ATGAACAAAAGCCAACTTTATCCGGATTCACCACTGACGGATCAGGACTTCAACCAATTAGACCAAAC
CGTGATTGAGGCTGCTCGTCGTCAGCTGGTGGGTCGTCGCTTCATTGAGTTATATGGCCCATTGGGGC
GTGGCATGCAGAGTGTCTTCAACGATATCTTCATGGAGTCTCATGAAGCGAAAATGGACTTCCAGGGC
AGCTTTGACACGGAGGTAGAGTCCTCCCGTCGTGTAAACTATACCATTCCGATGTTATATAAAGACTT
CGTGCTTTACTGGCGCGATCTGGAACAGAGCAAGGCACTCGATATTCCGATCGACTTTTCAGTGGCAG
CGAACGCTGCCCGCGACGTTGCGTTCCTGGAAGATCAGATGATTTTCCATGGAAGCAAAGAATTTGAT
ATCCCGGGTCTGATGAACGTGAAAGGTCGCCTGACCCATCTGATTGGCAATTGGTATGAGTCGGGTAA
CGCCTTTCAGGATATTGTGGAGGCCCGCAATAAATTACTCGAAATGAACCACAATGGCCCATATGCTC
TCGTGCTGTCCCCGGAGCTGTACTCACTCTTACATCGTGTGCATAAAGACACGAATGTGCTGGAGATC
GAACACGTGCGCGAGTTGATTACTGCTGGGGTTTTTCAGTCGCCTGTCCTCAAAGGGAAAAGTGGTGT
GATCGTAAACACCGGTCGCAACAATCTGGATTTGGCTATCTCGGAAGATTTTGAGACTGCATACCTGG
GCGAGGAAGGTATGAACCATCCCTTTCGCGTGTACGAGACAGTTGTTCTGCGCATCAAACGCCCGGCG
GCCATTTGTACTTTAATCGATCCGGAAGAATAA

#### 1.2 Codon-optimised sequence of LanM-QtEnc fusion gene

The following gene sequence shows the hexahistidine tag in underlined purple, LanM in bold green, TEV protease cleavage site in underlined red, and the QtEnc gene in bold blue.

The synthetic gene was inserted into a pETDuet-1 vector between NcoI and KpnI restriction sites by Gibson assembly.

ATGCATCACCACCATCATCACGGAGGAAGT**CCGACAACGACCACCAAAGTTGACATCGCCGCATTTGA
TCCAGATAAAGACGGAACGATCGACTTGAAAGAGGCACTGGCTGCTGGTTCGGCGGCATTTGATAAAC
TGGACCCGGATAAGGACGGTACTCTTGATGCAAAAGAATTAAAAGGGCGTGTCTCCGAAGCTGACTTG
AAGAAACTGGACCCTGACAACGACGGCACATTGGACAAGAAGGAGTACCTTGCAGCTGTGGAGCAATT
CAAAGCAGCGAACCCAGACAACGATGGCACGATTGACGCTCGCGAACTGGCTTCCCCGGCAGGTTCCG
CCTTGGTGAATCTTATTCGG**GGGGGCTCCGGTGGCTCGGAGAATCTTTATTTTCAGTCT**AACAAAAGC
CAACTTTATCCGGATTCACCACTGACGGATCAGGACTTCAACCAATTAGACCAAACCGTGATTGAGGC
TGCTCGTCGTCAGCTGGTGGGTCGTCGCTTCATTGAGTTATATGGCCCATTGGGGCGTGGCATGCAGA
GTGTCTTCAACGATATCTTCATGGAGTCTCATGAAGCGAAAATGGACTTCCAGGGCAGCTTTGACACG
GAGGTAGAGTCCTCCCGTCGTGTAAACTATACCATTCCGATGTTATATAAAGACTTCGTGCTTTACTG
GCGCGATCTGGAACAGAGCAAGGCACTCGATATTCCGATCGACTTTTCAGTGGCAGCGAACGCTGCCC
GCGACGTTGCGTTCCTGGAAGATCAGATGATTTTCCATGGAAGCAAAGAATTTGATATCCCGGGTCTG
ATGAACGTGAAAGGTCGCCTGACCCATCTGATTGGCAATTGGTATGAGTCGGGTAACGCCTTTCAGGA
TATTGTGGAGGCCCGCAATAAATTACTCGAAATGAACCACAATGGCCCATATGCTCTCGTGCTGTCCC
CGGAGCTGTACTCACTCTTACATCGTGTGCATAAAGACACGAATGTGCTGGAGATCGAACACGTGCGC
GAGTTGATTACTGCTGGGGTTTTTCAGTCGCCTGTCCTCAAAGGGAAAAGTGGTGTGATCGTAAACAC
CGGTCGCAACAATCTGGATTTGGCTATCTCGGAAGATTTTGAGACTGCATACCTGGGCGAGGAAGGTA
TGAACCATCCCTTTCGCGTGTACGAGACAGTTGTTCTGCGCATCAAACGCCCGGCGGCCATTTGTACT
TTAATCGATCCGGAAGAA**TAA

### 1.3 Codon-optimised sequence of mNeonGreen with targeting peptides for encapsulation

mNeonGreen expression constructs (gene in bold text) were designed with an *N*-terminal hexahistidine tag (underlined in purple), and a *C*-terminal targeting peptide sequence from either *Q. thermotolerans* or *T. maritima* (underlined in orange), using the second cassette of pCDFDuet-1 plasmid (NdeI and PacI restriction sites).

mNeon-Qt

ATGCACCACCATCATCATCACGGAGGTTCA**GTCTCCAAGGGAGAGGAGGACAATATGGCTAGTCTTCC**
**GGCCACTCATGAGTTACATATCTTCGGATCCATAAACGGCGTTGATTTCGATATGGTGGGTCAAGGCA**
**CTGGTAACCCCAATGATGGCTACGAAGAACTTAACTTAAAATCTACTAAAGGCGATCTTCAGTTTTCC**
**CCATGGATACTTGTGCCTCATATTGGCTACGGGTTTCACCAATATCTGCCTTATCCGGATGGAATGTC**
**CCCCTTCCAAGCCGCAATGGTAGACGGCAGTGGCTATCAAGTCCACCGTACCATGCAGTTTGAAGATG**
**GCGCATCCCTGACAGTTAATTATCGGTATACATACGAGGGGTCGCATATTAAAGGAGAGGCGCAAGTC**
**AAGGGGACAGGGTTTCCCGCCGATGGGCCAGTCATGACAAACTCGTTAACTGCCGCCGACTGGTGCAG**
**ATCGAAGAAAACCTACCCAAACGATAAGACGATCATATCTACCTTTAAATGGTCTTACACTACGGGTA**
**ACGGAAAACGCTACAGATCAACCGCGCGGACAACGTACACCTTTGCTAAGCCCATGGCAGCGAACTAC**
**TTGAAGAATCAGCCGATGTACGTGTTTAGAAAGACCGAGCTTAAACACTCGAAAACTGAATTGAATTT**
**TAAAGAATGGCAGAAAGCTTTTACGGACGTAATGGGCATGGACGAACTGTATAAG**TCCGGGTCGGGAG
GCTCCGGAAAAAAGAAAGGCTTTACTGTCGGGTCGTTAATTCAGTAA

mNeon-Tm

ATGCACCACCATCATCATCACGGAGGTTCA**GTCTCCAAGGGAGAGGAGGACAATATGGCTAGTCTTCC**
**GGCCACTCATGAGTTACATATCTTCGGATCCATAAACGGCGTTGATTTCGATATGGTGGGTCAAGGCA**
**CTGGTAACCCCAATGATGGCTACGAAGAACTTAACTTAAAATCTACTAAAGGCGATCTTCAGTTTTCC**
**CCATGGATACTTGTGCCTCATATTGGCTACGGGTTTCACCAATATCTGCCTTATCCGGATGGAATGTC**
**CCCCTTCCAAGCCGCAATGGTAGACGGCAGTGGCTATCAAGTCCACCGTACCATGCAGTTTGAAGATG**
**GCGCATCCCTGACAGTTAATTATCGGTATACATACGAGGGGTCGCATATTAAAGGAGAGGCGCAAGTC**
**AAGGGGACAGGGTTTCCCGCCGATGGGCCAGTCATGACAAACTCGTTAACTGCCGCCGACTGGTGCAG**
**ATCGAAGAAAACCTACCCAAACGATAAGACGATCATATCTACCTTTAAATGGTCTTACACTACGGGTA**
**ACGGAAAACGCTACAGATCAACCGCGCGGACAACGTACACCTTTGCTAAGCCCATGGCAGCGAACTAC**
**TTGAAGAATCAGCCGATGTACGTGTTTAGAAAGACCGAGCTTAAACACTCGAAAACTGAATTGAATTT**
**TAAAGAATGGCAGAAAGCTTTTACGGACGTAATGGGCATGGACGAACTGTATAAG**TCAGGCGGGAACA
CAGGAGGCGATTTAGGCATTCGCAAGTTATAA

### 1.4 Codon-optimised sequence of TEV protease

The TEV protease S219V mutant expression construct (gene in bold blue) was designed with an *N*-terminal hexahistidine tag (underlined in purple) followed by a dihydrolipoyllysine-binding domain for solubility (bold green), inserting into a pETDuet-1 vector between the NcoI and EcoRI restriction sites.

ATGCACCACCACCACCACCACAGCGGC**GCTTTTGAATTTAAGCTGCCGGACATTGGCGAAGGCATCCA**
**CGAAGGTGAAATTGTCAAATGGTTTGTGAAACCGGGCGATGAAGTGAACGAAGACGATGTATTGTGCG**
**AAGTGCAAAATGACAAGGCGGTTGTCGAAATTCCCTCCCCGGTCAAAGGGAAAGTGCTTGAAATCCTC**
**GTCCCGGAGGGAACAGTGGCAACGGTCGGGCAAACGCTCATCACGCTCGATGCGCCGGGTTATGAAAA**
**CATGACG**AGCAGCGGCCTGGTGCCGCGCGGATCC**GGAGAAAGCTTGTTTAAGGGACCACGTGATTACA**
**ACCCGATATCGAGCACCATTTGTCATTTGACGAATGAATCTGATGGGCACACAACATCGTTGTATGGT**
**ATTGGATTTGGTCCCTTCATCATTACAAACAAGCACTTGTTTAGAAGAAATAATGGAACACTGTTGGT**
**CCAATCACTACATGGTGTATTCAAGGTCAAGAACACCACGACTTTGCAACAACACCTCATTGATGGGA**
**GGGACATGATAATTATTCGCATGCCTAAGGATTTCCCACCATTTCCTCAAAAGCTGAAATTTAGAGAG**
**CCACAAAGGGAAGAGCGCATATGTCTTGTGACAACCAACTTCCAAACTAAGAGCATGTCTAGCATGGT**

**GTCAGACACTAGTTGCACATTCCCTTCATCTGATGGCATATTCTGGAAGCATTGGATTCAAACCAAGG**
**ATGGGCAGTGTGGCAGTCCATTAGTATCAACTAGAGATGGGTTCATTGTTGGTATACACTCAGCATCG**
**AATTTCACCAACACAAACAATTATTTCACAAGCGTGCCGAAAAACTTCATGGAATTGTTGACAAATCA**
**GGAGGCGCAGCAGTGGGTTAGTGGTTGGCGATTAAATGCTGACTCAGTATTGTGGGGGGGCCATAAAG**
**TTTTCATGGTCAAACCTGAAGAGCCTTTTCAGCCAGTTAAGGAAGCG**AATAGGGCTCATGAATGA

### 2. Protein sequences

| Name | Protein Sequence |
| --- | --- |
| Wt QtEnc | MNKSQLYPDSPLTDQDFNQLDQTVIEAARRQLVGRRFIELYGPLGRGMQ<br>SVFNDIFMESHEAKMDFQGSFDTEVESSRRVNYTIPMLYKDFVLYWRDL<br>EQSKALDIPIDFSVAANAARDVAFLEDQMIFHGSKEFDIPGLMNVKGRLT<br>HLIGNWYESGNAFQDIVEARNKLLEMNHNGPYALVLSPELYSLLHRVHK<br>DTNVLEIEHVRELITAGVFQSPVLKGKSGVIVNTGRNNLDLAISEDFETA<br>YLGEEGMNHPFRVYETVVLRIKRPAAICTLIDPEE* |
| 6xHis-LanM-TEV-QtEnc | MHHHHHHGGSPTTTTKVDIAAFDPDKDGTIDLKEALAAGSAAFDKLDP<br>DKDGTLDAKELKGRVSEADLKKLDPDNDGTLDKKEYLAAVEQFKAAN<br>PDNDGTIDARELASPAGSALVNLIRGGSGGSENLYFQSNKSQLYPDSPLT<br>DQDFNQLDQTVIEAARRQLVGRRFIELYGPLGRGMQSVFNDIFMESHEA<br>KMDFQGSFDTEVESSRRVNYTIPMLYKDFVLYWRDLEQSKALDIPIDFSV<br>AANAARDVAFLEDQMIFHGSKEFDIPGLMNVKGRLTHLIGNWYESGNA<br>FQDIVEARNKLLEMNHNGPYALVLSPELYSLLHRVHKDTNVLEIEHVRE<br>LITAGVFQSPVLKGKSGVIVNTGRNNLDLAISEDFETAYLGEEGMNHPFR<br>VYETVVLRIKRPAAICTLIDPEE* |
| 6xHis-mNeon-Qt CLP | MHHHHHHGGSVSKGEEDNMASLPATHELHIFGSINGVDFDMVGQGTGN<br>PNDGYEELNLKSTKGDLQFSPWILVPHIGYGFHQYLPYPDGMSPFQAAM<br>VDGSGYQVHRTMQFEDGASLTVNYRYTYEGSHIKGEAQVKGTGFPADG<br>PVMTNSLTAADWCRSKKTYPNDKTIISTFKWSYTTGNGKRYRSTARTTY<br>TFAKPMAANYLKNQPMYVFRKTELKHSKTELNFKEWQKAFTDVMGM<br>DELYKSGSGGSGKKKGFTVGSLIQ* |
| 6xHis-mNeon-Tm CLP | MHHHHHHGGSVSKGEEDNMASLPATHELHIFGSINGVDFDMVGQGTGN<br>PNDGYEELNLKSTKGDLQFSPWILVPHIGYGFHQYLPYPDGMSPFQAAM<br>VDGSGYQVHRTMQFEDGASLTVNYRYTYEGSHIKGEAQVKGTGFPADG<br>PVMTNSLTAADWCRSKKTYPNDKTIISTFKWSYTTGNGKRYRSTARTTY<br>TFAKPMAANYLKNQPMYVFRKTELKHSKTELNFKEWQKAFTDVMGM<br>DELYKSGGNTGGDLGIRKL* |
| 6xHis-Lipoyl domain-TEV Protease | MHHHHHHSGAFEFKLPDIGEGIHEGEIVKWFVKPGDEVNEDDVLCEVQ<br>NDKAVVEIPSPVKGKVLEILVPEGTVATVGQTLITLDAPGYENMTSSGLV<br>PRGSGESLFKGPRDYNPISSTICHLTNESDGHTTSLYGIGFGPFIITNKHLF<br>RRNNGTLLVQSLHGVFKVKNTTTLQQHLIDGRDMIIIRMPKDFPPFPQKL<br>KFREPQREERICLVTTNFQTKSMSSMVSDTSCTFPSSDGIFWKHWIQTKD<br>GQCGSPLVSTRDGFIVGIHSASNFTNTNNYFTSVPKNFMELLTNQEAQQ<br>WVSGWRLNADSVLWGGHKVFMVKPEEPFQPVKEANRAHE* |

### 3. Supplementary SEC data, PAGE gels and TEM images

#### 3.1 Additional data for the purification of LanM-QtEnc

SDS-PAGE was conducted on Any kD Mini-Protean TGX Stain-Free protein gels (Bio-Rad) according to manufacturer protocols.

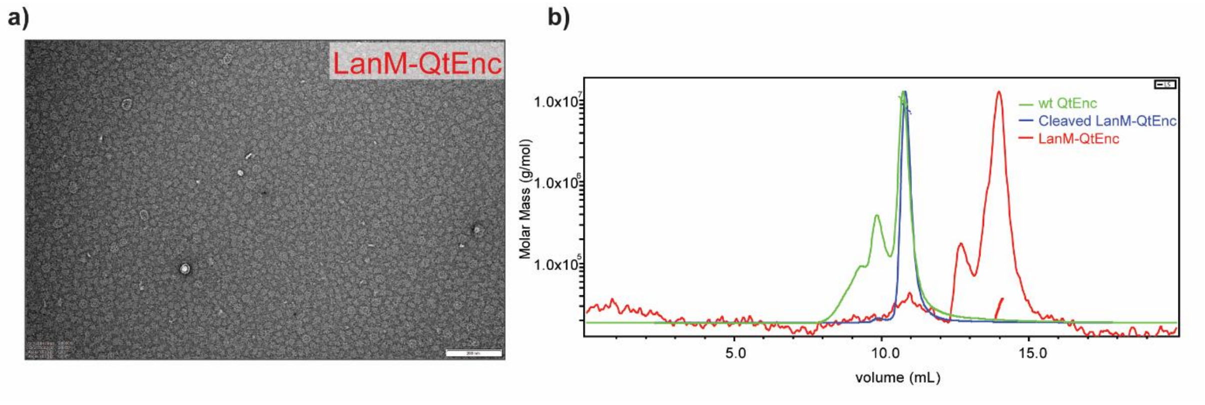


**Figure S3.1. a)** Negative stain transmission electron microscopy of LanM-QtEnc fusion demonstrates mostly unassembled proteins. Images were taken at 100000× magnification (scale bar = 200 nm). **b)** Size exclusion chromatography with light scattering detection on Bio SEC-5 2000 Å HPLC column shows wt QtEnc (green solid line) and *in vitro* assembled cleaved LanM-QtEnc (blue solid line) have similar elution profiles with their molar mass calculated from differential refractive index measurement as 9.8 ± 0.2 MDa and 8.2 ± 0.2 MDa respectively. LanM-QtEnc (red solid line) fusion elutes much later and its molar mass calculated as 28.4 kDa ± 0.8 kDa. Molar mass is represented on each elution profile as dotted line of same colour.

#### 3.2 Additional native PAGE gel images

The following uncropped gels contain the BN-PAGE data shown Figures 2b, 3c, 4c and 4d of the main manuscript.

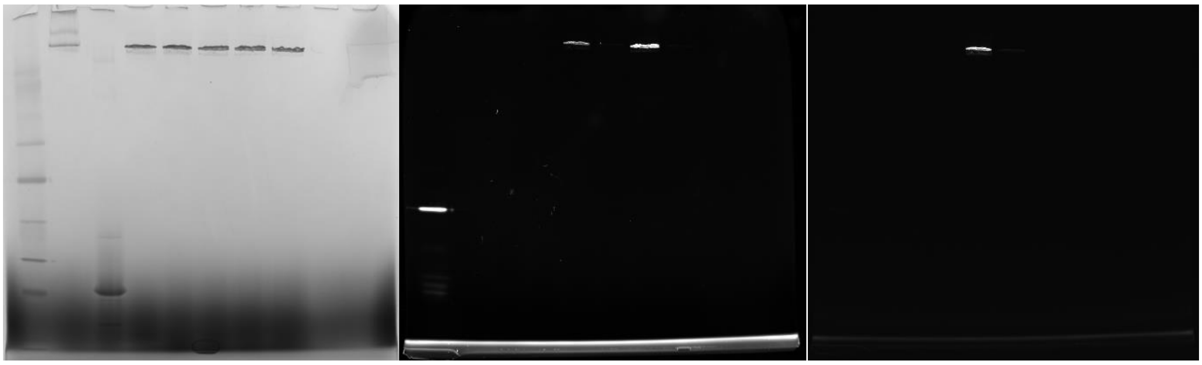

**Figure S3.2. BN-PAGE analysis of encapsulin samples.** Samples loaded from left to right are NativeMark ladder, wt QtEnc, LanM-QtEnc, cleaved LanM-QtEnc, loading of mNeon-Qt, mNeon-Tm, TMR-Qt, TMR-Tm. **Left:** Coomassie-stained BN-PAGE gel. **Middle:** 302 nm excitation and 590/110 nm emission filter for TAMRA. **Right:** 460-490 nm excitation and 532/28 nm emission filter for mNeonGreen.

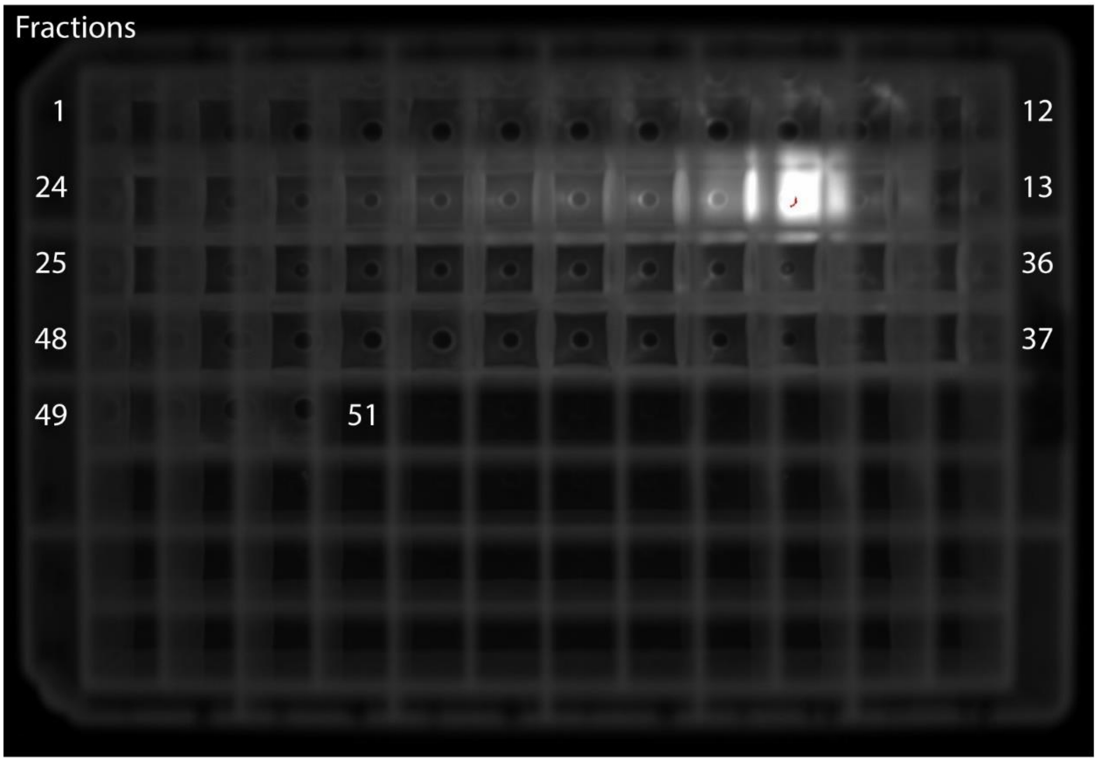


**Figure S3.3.** Fractions from size-exclusion chromatography of mNeon-Qt *in vitro* packaged into encapsulin cages, collected in serpentine order in deep-well plates and visualised by fluorescence emission in a ChemiDoc MP imager with the Alexa488 program settings (460-490 nm excitation, 532/28 nm emission filter). The encapsulin peak (fraction 15) is the only fraction that exhibits fluorescence.

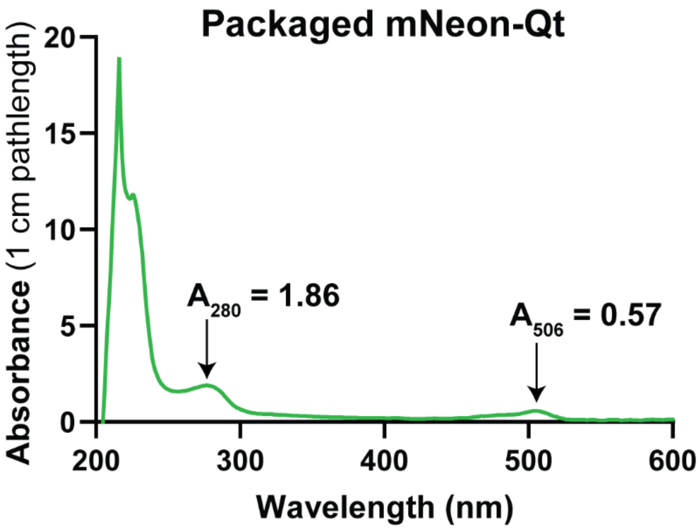


**Extinction coefficients:**

$QtEnc_{\varepsilon 280}$ = 25900 $M^{-1}$ $cm^{-1}$ (ProtParam)

$mNeon\text{-}Qt_{\varepsilon 506}$ = 116000 $M^{-1}$ $cm^{-1}$ (Shaner *et al., Nat. Methods* 2013)

$mNeon\text{-}Qt_{A280}/mNeon\text{-}Qt_{A506}$ = 0.537 (from measured absorbance of mNeon-Qt alone)

**Calculations:**

**[mNeon-Qt]** = $Total_{A506}/mNeon\text{-}Qt_{\varepsilon 506}$ = 0.57/116000 = 5 × $10^{-6}$ M = **5 µM**

**[QtEnc]** = $Qt\text{-}Enc_{A280}/QtEnc_{\varepsilon 280}$

= ($Total_{A280}$ – $mNeon\text{-}Qt_{A280}$)/$QtEnc_{\varepsilon 280}$

= ($Total_{A280}$ – (0.537 × $mNeon\text{-}Qt_{A506}$))/$QtEnc_{\varepsilon 280}$

= (1.86 – (0.537 × 0.57))/25900 = 60 × $10^{-6}$ M = **60 µM**

**[mNeon-Qt]:[QtEnc] = 1:12 packaged ratio**

**Figure S3.4**. Packaging of mNeon-Qt within *in vitro* assembled QtEnc is highly efficient**.**

**Top:** UV Vis absorbance spectrum of the encapsulin peak from Superose 6-Increase separation collected on a Nanodrop ND-1000 spectrophotometer. Arrows indicated the protein absorbance at 280 nm and the mNeon-specific absorbance at 506 nm.

**Bottom:** Calculated concentration of each protein species in the packaged sample after purification.

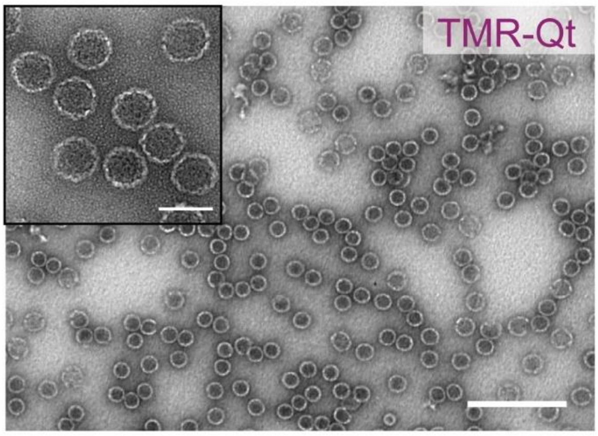


**Figure S3.5.** Negative stain transmission electron microscopy of *in vitro* packaged TMR-Qt into QtEnc cages have similar morphology and size to wild-type QtEnc. The larger images were taken at 40000× magnification (scale bar = 250 nm), while the inset images were taken at 200000× magnification (scale bar = 50 nm).

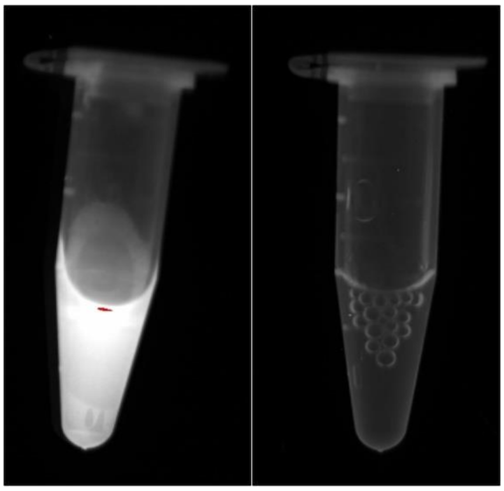

**Figure S3.6.** The purified encapsulin fractions from size-exclusion chromatography of TMR-Qt (left) and TMR-Tm (right) when *in vitro* packaged into encapsulin cages, visualised by fluorescence emission in a ChemiDoc MP imager with the Alexa488 program settings (460-490 nm excitation, 532/28 nm emission filter). Only TMR-Qt exhibits fluorescence associated with successful packaging of the synthetic dye.

### 4. Peptide synthesis and characterisation

**Table S4.1. Summary of peptide sequences and mass spectrometry data.**

| Name | Sequence | Expected mass | Observed mass |
|---|---|---|---|
| TMR-Tm | TMR-SENTGGDLGIRKL-$NH_2$ | $[M + 2H]^{2+}$ = 886.5 | $[M + 2H]^{2+}$ = 886.4 |
| TMR-Qt | TMR-KKKGFTVGSLIQ-$NH_2$ | $[M + 2H]^{2+}$ = 859.5 | $[M + 2H]^{2+}$ = 859.5 |
| Aldox-Qt | Ac-C(Aldox)KKKGFTVGSLIQ-$NH_2$ | $[M + 2H]^{2+}$ = 1101.1 | $[M + 2H]^{2+}$ = 1101.1 |

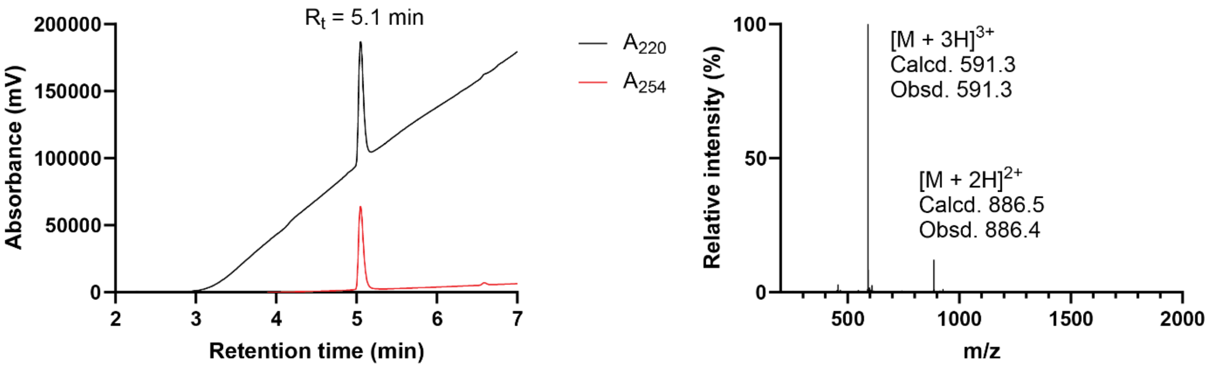


**Figure S4.2. Characterisation of TMR-Tm by LCMS.** UV absorbance was monitored at 220 and 254 nm, while mass spectra were obtained in electrospray ionisation (ESI) mode.

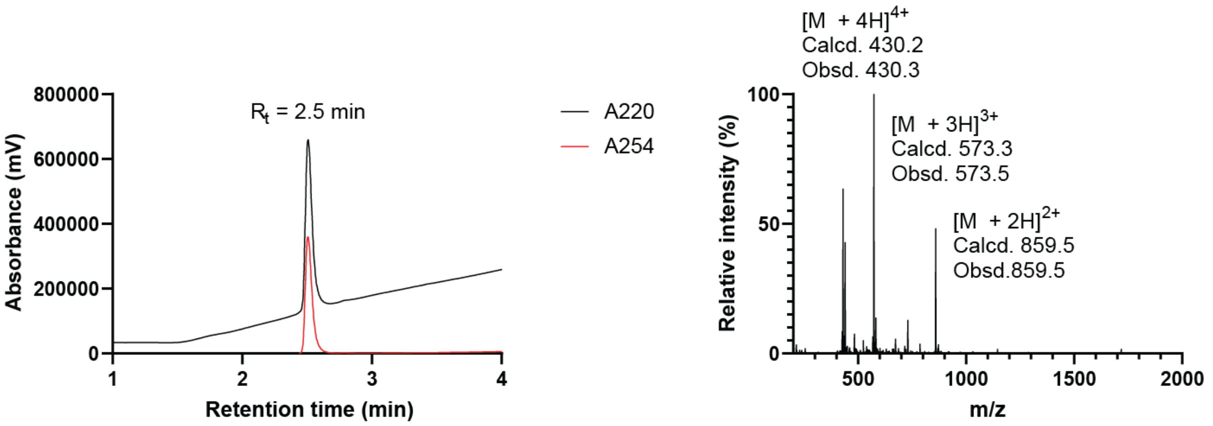


**Figure S4.3. Characterisation of TMR-Qt by LCMS.** UV absorbance was monitored at 220 nm and 254 nm, while mass spectra were obtained in electrospray ionisation (ESI) mode.

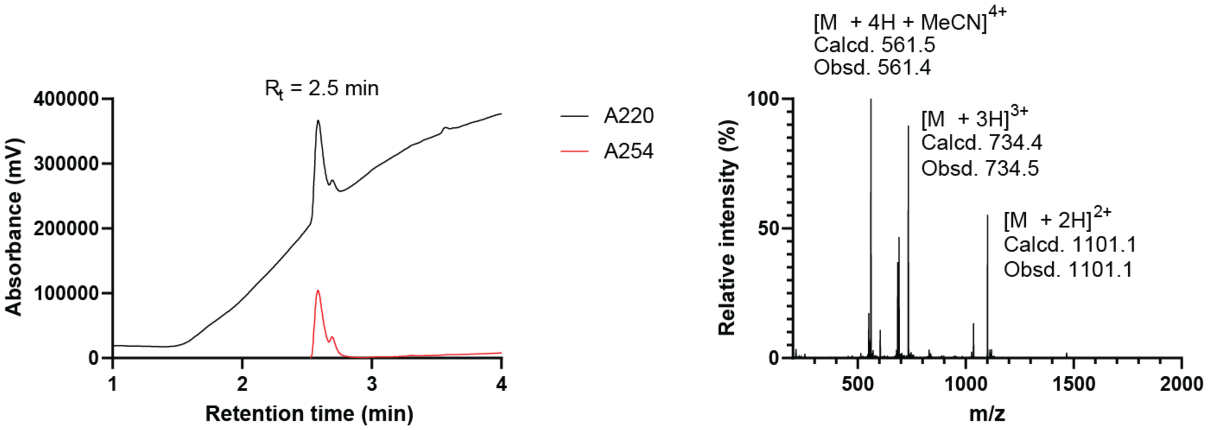


**Figure S4.4. Characterisation of Aldox-Qt by LCMS.** UV absorbance was monitored at 220 nm and 254 nm, while mass spectra were obtained in electrospray ionisation (ESI) mode.
